# CRISPRedict: The case for simple and interpretable efficiency prediction for CRISPR-Cas9 gene editing

**DOI:** 10.1101/2022.04.07.486362

**Authors:** Vasileios Konstantakos, Anastasios Nentidis, Anastasia Krithara, Georgios Paliouras

## Abstract

The development of the CRISPR-Cas9 technology has provided a simple yet powerful system for targeted genome editing. Compared with previous gene-editing tools, the CRISPR-Cas9 system identifies target sites by the complementarity between the guide RNA (gRNA) and the DNA sequence, which is less expensive and time-consuming, as well as more precise and scalable. To effectively apply the CRISPR-Cas9 system, researchers need to identify target sites that can be cleaved efficiently and for which the candidate gRNAs have little or no cleavage at other genomic locations. For this reason, numerous computational approaches have been developed to predict cleavage efficiency and exclude undesirable targets. However, current design tools cannot robustly predict experimental success as prediction accuracy depends on the assumptions of the underlying model and how closely the experimental setup matches the training data. Moreover, the most successful tools implement complex machine learning and deep learning models, leading to predictions that are not easily interpretable.

Here, we introduce *CRISPRedict*, a simple linear model that provides accurate and interpretable predictions for guide design. Comprehensive evaluation on twelve independent datasets demonstrated that *CRISPRedict* has an equivalent performance with the currently most accurate tools and outperforms the remaining ones. Moreover, it has the most robust performance for both U6 and T7 data, illustrating its applicability to tasks under different conditions. Therefore, our system can assist researchers in the gRNA design process by providing accurate and explainable predictions. These predictions can then be used to guide genome editing experiments and make plausible hypotheses for further investigation. The source code of *CRISPRedict* along with instructions for use is available at https://github.com/VKonstantakos/CRISPRedict.

## 1 Introduction

The Clustered Regularly Interspaced Short Palindromic Repeat (CRISPR)/ CRISPR-associated protein 9 (Cas9) technology, a prokaryotic immune system [1–5], has been rapidly and widely adopted by the scientific community to target and modify the genomes of a vast array of cells and organisms [6, 7]. It includes two key components: a synthetic single-guide RNA (sgRNA) and the Cas9 nuclease [7]. The sgRNA is an engineered sequence that combines all the components of the naturally occurring guide RNA complex. It consists of the native CRISPR RNA (crRNA) that directs Cas9 to the corresponding target site, and a trans-activating crRNA (tracrRNA) which forms a scaffold for Cas9 binding [8]. Thus, precise targeting can be achieved by synthesizing an sgRNA with a guide domain (gRNA) complementary to the target sequence while keeping the tracrRNA part constant. In the engineered CRISPR-Cas9 system, this gRNA is a 20-nucleotide (nt) sequence at the 5^t^ end of the sgRNA and is analogous to crRNA in the prokaryotic system [7].

Similar to the natural system, the gRNA guides Cas9 to cleave the DNA at a specific genomic locus based on sequence match, resulting in a double-stranded DNA break. This break occurs precisely 3 nt upstream of a protospacer adjacent motif (PAM) sequence [7, 9]. For CRISPR-Cas9, the PAM sequence is 5^t^-NGG-3^t^ (i.e., any nucleotide followed by two guanines). Therefore, target recognition requires both base pairing to the gRNA sequence and the presence of the PAM adjacent to the targeted sequence [7]. Once bound to the target sequence, the HNH and RuvC nuclease domains of Cas9 will cleave the DNA strands complementary and non-complementary to the guide sequence, leaving a blunt-ended DNA double-strand break (DSB) [7]. Following the break, random insertions or deletions (indels) can be generated via the non-homologous end-joining (NHEJ) pathway, leading to frameshift mutations and functional gene knockout. Alternatively, a desired modification can be introduced through homology-directed repair (HDR) when provided with a DNA template (gene knock-in) [10, 11].

To effectively apply the CRISPR-Cas9 system for gene editing, researchers need to identify target sites that can be cleaved efficiently and for which the candidate gRNAs have little or no cleavage at other genomic locations. Since multiple target sites can be available for a specific application, identifying the one with maximal on-target activity and minimal off-target effects is a significant challenge. However, variable activities among different gRNAs still represent a limitation, leading to inconsistent target efficiency. In addition, the specific features that determine on-target activity remain largely unexplored.

Therefore, accurate prediction of gRNA activity could facilitate the design process by excluding undesirable targets based on predicted low efficiency [12]. For this reason, numerous computational tools for guide design have been developed. The existing tools fall into three classes, namely alignment-based, hypothesis-driven, and learning-based methods [12]. Alignment-based tools retrieve suitable gRNAs from the given genome by locating the PAM sequence. For instance, *CRISPRdirect* [13] performs gRNA selection by investigating the entire genome for perfect matches with the candidate target sequence and the seed sequence flanking the PAM. Hypothesis-driven tools score the guide sequences according to empirically derived, handcrafted rules (e.g., GC content). For example, CHOPCHOP [14] initially used two metrics to score a given gRNA; its GC content - ideally between 40% and 80% - and whether the gRNA contains a G at position 20. Learning-based tools predict the gRNA efficiency using models that are trained on datasets of CRISPR experiments and incorporate various features. For example, Azimuth 2.0 [15], a gradient-boosted regression tree (GBRT) model, includes sequence features, thermodynamic features, and the location of the target within a gene to evaluate the efficiency of candidate gRNAs.

Reported results suggest that the latter two types of tools perform better than the alignment-based ones because they take into account many different features [16]. For learning-based tools, in particular, these features are combined by models that are generated through machine learning [12, 17]. Thus, they seem to perform better than alignment-based and hypothesis-driven tools [18]. Moreover, a shift has been observed recently in the learning-based category. While the initial learning-based tools relied on conventional machine learning methods, several deep learning-based methods have been explored lately for gRNA activity prediction [19–23].

However, the accuracy of machine learning-based tools varies widely among the constructed features and the testing datasets [18, 24]. Moreover, the hand-crafted features may result in redundancy, leading to inaccurate and non-interpretable predictions. For example, two state-of-the-art tools, Azimuth 2.0 [15] and sgDesigner [25], include 582 and 302 features, respectively, making their predictions hard to explain. Similarly, the recent deep learning tools also lack interpretability. Despite their ability to automatically identify important features, it is unclear how they derive a certain prediction. Furthermore, complex models are not necessarily better than simpler ones. Previous studies have, in fact, shown that the right method may differ per task and conventional machine learning tools can outperform the deep learning ones [18, 26]. Therefore, choosing the proper algorithm depends on the task and may not be based solely on predictive accuracy. In particular, interpretability, the ability to identify important features, and their interactions are all key factors that can influence the final choice.

In this work, we introduce *CRISPRedict*, a simple and interpretable linear model that predicts guide activities with comparable performance to the current state-of-the-art tools. The key goal of our system is to assist researchers in the gRNA design process by providing accurate and explainable predictions. These predictions can then be used to guide genome editing experiments and make plausible hypotheses for further investigation.

## 2 Materials and Methods

### 2.1 Data collection and processing

This section describes the process we followed to collect, extract and prepare the data used in our study. In total, we gathered 14 publicly available datasets that have been previously used to train and evaluate gRNA design tools. Two of those were used to implement *CRISPRedict*, while the remaining ones were used for testing purposes. Previous studies [22, 24, 27] have shown that the type of promoter for gRNA expression (i.e., U6 or T7) strongly influences the suitability and performance of the prediction model. Therefore, we created two versions of *CRISPRedict* to reflect the different experimental conditions in the guide design process. The training datasets were chosen based on their size, homogeneity, and previous evaluation results [18, 21, 28]. Similarly, to compare the performance of *CRISPRedict* with the state-of-the-art methods on datasets of different expression systems, we collected 12 datasets from independent studies. These studies reflect the different conditions and tasks that can arise during the use of *CRISPRedict*. Thus, they allow for a comprehensive evaluation of the two model variants. The extracted datasets with their respective studies are the following:

- Training datasets

- Kim (*DeepSpCas9* training dataset) [21].
- Moreno-Mateos [24].

- Testing datasets

- U6 Regression: Labuhn [19,29], Shalem [30], Koike-Yusa [31], Kim (*DeepSpCas9* testing dataset) [21].
- T7 Regression: Shkumatava [24], Gagnon [24], Varshney [24], Teboul [24].
- Classification: Koike-Yusa [31], Wang [31], Chari [32], Hiranniramol (*sgDesigner* training dataset) [25].

Some of the extracted datasets represented cleavage efficiency on a different scale, based on the technique used to measure the actual outcome. For this reason, we rescaled the knockout efficiencies to obtain a standardized efficiency measurement that can be used for training and evaluation. In particular, we applied a Min-Max rescaling procedure to the cleavage efficiency of the dataset. The Min-Max normalization maps a value in the range [0, 1] via the following equation:

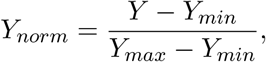

where *Y*_*min*_ and *Y*_*max*_ are, respectively, the minimum and maximum efficiency value of the dataset, *Y* is the original efficiency and *Y*_*norm*_ is the normalized value.

In addition, the datasets included sequences of variable length, while we needed 23-nt and 30-nt sequences for model training and evaluation. Therefore, we extracted sequences in the appropriate form using the provided sequence context. For example, in the case of the Moreno-Mateos dataset [24, 28], the 23-nt sequences were found in the long 100-nt sequences and then extracted along with the four preceding and the three following nucleotides, obtaining 30-nt sequences.

#### 2.1.1 Training datasets

We used two public experimentally-validated efficiency datasets to implement *CRISPRedict*. In particular, the training datasets of *DeepSpCas9* (“Kim”) [21] and *CRISPRscan* (“Moreno-Mateos”) [24,28] were used to create two models suitable for gRNAs expressed under the U6 and T7 promoter, respectively. This choice was based on previous studies that showed that the accuracy of a prediction model strongly depends on whether the gRNA is expressed from a U6 promoter in cells or from a T7 promoter in vitro [22, 24, 27].

The Kim dataset was gathered from the corresponding study [21] and was used without any changes. The Moreno-Mateos dataset was extracted from Haeussler et al. [24]. After selecting the necessary sequences, we rescaled the knockout efficiencies and extracted 30-nt sequences using the procedures described in Section 2.1.

#### 2.1.2 Testing datasets

A total of 12 testing datasets were gathered from published studies to evaluate the performance of our tool under three scenarios. These scenarios reflect the different experimental conditions and tasks that can arise during the model’s use. They consist of two regression schemes and a classification scheme. The regression scenarios include gRNAs transcribed from a U6 or a T7 promoter (U6 regression/T7 regression), while the classification one includes labeled gRNAs transcribed from a U6 promoter.

Regarding the U6 regression scheme, we gathered four datasets, namely Labuhn [19,29], Shalem [30], Koike-Yusa [31] and Kim [21]. The Labuhn dataset [29] was gathered from the supplementary material provided by Chuai et al. [19]. This dataset utilized fluorescent reporter knockout assays with verification at selected endogenous loci for gRNA knockout efficiency measurement. It contains a total of 425 gRNAs for HEL cells [29]. The Shalem dataset was provided from Doench et al. [30] and contains 1278 gRNAs targeting 414 genes. The gRNA efficiency is expressed as the *log*_2_ fold change in abundance during two weeks of growth in A375 cells. The Koike-Yusa dataset was provided by Xu et al. [31]. It originally contained 87897 gRNAs targeting 19150 mouse protein-coding genes in mouse embryonic stem cells (mESCs) [33]. By posterior analysis, 311 essential genes were identified and 1064 gRNAs were retained [31]. The Kim test dataset was provided by the corresponding study [21] and includes 542 target sequences that were not included in the training dataset of *DeepSpCas9*.

For the T7 regression scenario, we extracted four datasets from Haeussler et al. [24]: Shkumatava, Gagnon, Varshney, and Teboul. All the datasets included guides that were transcribed - either in vitro (Shkumatava, Gagnon) or in vivo (Varshney, Teboul) - with the T7 promoter. In particular, the Shkumatava dataset is a set of 163 guide sequences from 11 different loci in zebrafish. The gRNA efficiency is the number of mutated sequencing clones obtained from zebrafish embryos. Similarly, the data from Gagnon include 111 target sequences from zebrafish, with the percentage of insertions-deletions (indels) representing the gRNA efficiency. On the other hand, the Varshney and Teboul dataset include 102 and 30 guides transcribed in vivo in zebrafish and mouse, respectively. They both use the percentage of mutated embryos to capture the actual gRNA efficiency.

Finally, for the classification evaluation scheme, we used four datasets which included gRNAs labeled as efficient or inefficient. The datasets were Koike-Yusa [31], Wang [31], Chari [32], and Hiranniramol [25]. The guides included in these datasets were transcribed using the U6 promoter in human (Wang, Chari, Hiranniramol) and mouse (Koike-Yusa) cell lines. The datasets Wang and Koike-Yusa are originally from Wang et al. [34] and Koike-Yusa et al. [33], but were used as processed by Xu et al. [31]. They include 2077 and 1064 gRNAs transcribed in human HL60 and mouse embryonic stem cells (mESC), respectively. The Chari dataset was directly retrieved from the study’s supplementary material [32] and extracted from Haeussler et al. [24] for reproducibility. Similar to the approach of Chari et al. [32] and Haeussler et al. [24], we used only the SpCas9 dataset, which includes 279 gRNAs transcribed in human 293T cells. The Hiranniramol dataset was obtained from the corresponding study [25] and was originally used to train *sgDesigner*. It is a plasmid library expressed in human 293T cells, which includes 746 functional and 563 nonfunctional gRNAs transcribed from a U6 promoter.

We extracted 23-nt and 30-nt sequences and rescaled the efficiency measurements from all the testing datasets using the methods described in Section 2.1. Python scripts to reproduce our results are available in the provided GitHub page.

### 2.2 Development of CRISPRedict

We created a U6 and a T7 variant of *CRISPRedict* to improve the prediction ability of our model under different experimental conditions. Each variant was trained on a different dataset as described in Section 2.1.1, using a similar modeling pipeline. To construct the final model for our task, we combined an algorithm comparison and a feature selection strategy.

First, we defined the initial feature set using sequence characteristics that have been shown to influence cleavage efficiency [15, 18, 19, 28–32, 34, 35]. This includes overall and position-specific nucleotide composition, as well as features that reflect the structural properties of the guide sequence. We then applied a multi-step feature selection strategy to arrive at a minimal and relevant-only feature subset. We also defined a set of different models to evaluate at each step and select the appropriate one for each use-case. Given the nature of the problem and the goal of our study, we chose the model with the highest performance and lowest complexity.

#### 2.2.1 Feature construction

We constructed features from target sequences using a similar approach to *Azimuth 2*.*0* [15] *and CRISPRpred* [36]. *In particular, we extended the sequences to 30 nucleotides (nt), in the form of N*_4_*N*_20_*NGGN*_3_, where *N*_*i*_ represents any *i* nucleotides. Thus, the first 4 nt and the last 3 nt were also extracted together with the original 20-nt spacer and the PAM NGG. We used the 30-nt sequences to create three groups of features: (i) nucleotide composition, (ii) position-specific nucleotides, and (iii) miscellaneous features. We denote the 30-nt sequence as S and adenine, guanine, thymine, cytosine as A, G, T, C, respectively.

Regarding the overall nucleotide composition, the number of each single nucleotide (e.g., how many A) in S was counted, resulting in 4 features. Similarly, we considered all dinucleotide or trinucleotide combinations and got 4^2^ = 16 and 4^3^ = 64 features, respectively.

For the position-specific nucleotides, we treated each position of a nucleotide as a binary value and created 30*∗* 4 = 120 features. For instance, we created 4 features for the first position of S based on the presence of any nucleotide (i.e., A, C, G, T). Similarly, we created another 29 *∗* 4^2^ = 464 binary features by checking the presence of all dinucleotide combinations at each position. For example, if the first nucleotide is A and the second nucleotide is C then AC 1 = 1, otherwise AC 1 = 0, meaning the absence of AC in the first and second positions. Moreover, we combined three nucleotides that are adjacent and repeated the previous feature construction process to create 28 *∗* 4^3^ = 1792 features.

Finally, we extracted 6 features that have been shown to influence gRNA efficiency, but do not fit into a single category. Specifically, the total GC content [29, 30, 34, 37], the number of As in the middle [30, 31, 34], and the presence of certain motifs (e.g., GGGG, TT, GCC) [35, 38, 39], which are known to strongly impact guide activity, were used for the initial feature set.

Thus, we extracted a total of 2466 features from the provided 30-nt sequences. Thermodynamic and structural features were also calculated but did not improve the final performance; thus, they were not included. All features were created using custom Python scripts.

#### 2.2.2 Feature selection

Given that the initial set of 2466 features was over-determined, we applied multiple steps of feature selection to remove any redundant and unnecessary features and choose the feature subset with the lowest generalization error.

First, we incorporated L1-based feature selection, as implemented in [40]. Since the L1-regularization forces coefficients for unnecessary features to be zero, we trained a model with the initial feature set and kept only the features with weights greater than 10^−6^.

Feature ranking with recursive feature elimination and cross-validated selection (RFECV) was then implemented by training a L1-regularized linear regression. During this procedure, the least important feature was pruned from the current set and the process was repeated until the optimal number of features was reached. This number was determined based on the model’s performance under 5-fold cross-validation. Spearman correlation was used as the evaluation measure for keeping the best features.

Based on the resulting features of the previous steps, our goal was to keep less than 50 features in order to gain interpretability without sacrificing accuracy. To accomplish that, we implemented an additional feature selection step based on Genetic Algorithms (GA). In particular, we used the Sklearn-genetic package [41] to determine the best 50, 40, 30, 20, and 10 features for the two model variants (i.e., U6 and T7). An L2-regularized linear regression model [40] was evaluated using Spearman correlation under 5-fold cross-validation for 500 generations. The feature sets with the best performance were saved to guide the final feature selection step. For more details about our GA implementation, the reader is referred to the provided source code and its documentation.

After manually inspecting the alternative feature sets produced by the GA feature selection step, we identified features that were present in most sets and also grouped some similar features into a single one. For example, we transformed the “TG 21”, “TG 22”, and “TG 23” features into one “TG end” feature, representing the presence of the dinucleotide TG in the last positions of the spacer sequence. This way, we managed to keep the most important features while limiting their number.

#### 2.2.3 Algorithm selection

We used the following models in our experiments: (i) Linear Regression (LR), (ii) Decision Tree (DT), (iii) Support Vector Regressor (SVR), (iv) Random Forest (RF), and (v) Extreme Gradient Boost (XGB). The chosen models were compared after each step of feature selection using 10-fold cross-validation. We selected these models because they differ in their complexity and their assumptions. Thus, they allow to capture various modelling aspects and determine which algorithm is more appropriate for our task. All models were implemented with the Scikit-learn [40] and XGBoost [42] packages in Python and used the default settings. The only exception was the SVR model, which used a radial basis function (RBF) kernel with the parameter gamma set to ‘auto’.

#### 2.2.4 Model construction

Following the feature selection strategy described in Section 2.2.2, we managed to greatly decrease the number of features without sacrificing accuracy. In particular, the L1-based selection reduced the initial 2466 features to 273 and 125 features for the U6 and T7 dataset, respectively. These features were then used for the RFECV procedure. The resulting scores with the corresponding number of features can be seen in Fig 1. We noticed that the model’s performance plateaued after the first 100 features and reached its maximum at 173 features for the U6 dataset. For the T7 dataset the plateau was observed at 70 features and the maximum at 93. Thus, we decreased the number of features to 173 and 93 for the U6 and T7 model, respectively, without affecting its performance. Finally, we further reduced the number of features based on the results of the GA and manual feature selection. The final feature sets included 28 and 25 features for the U6 and T7 variant, respectively.

**Figure 1:**
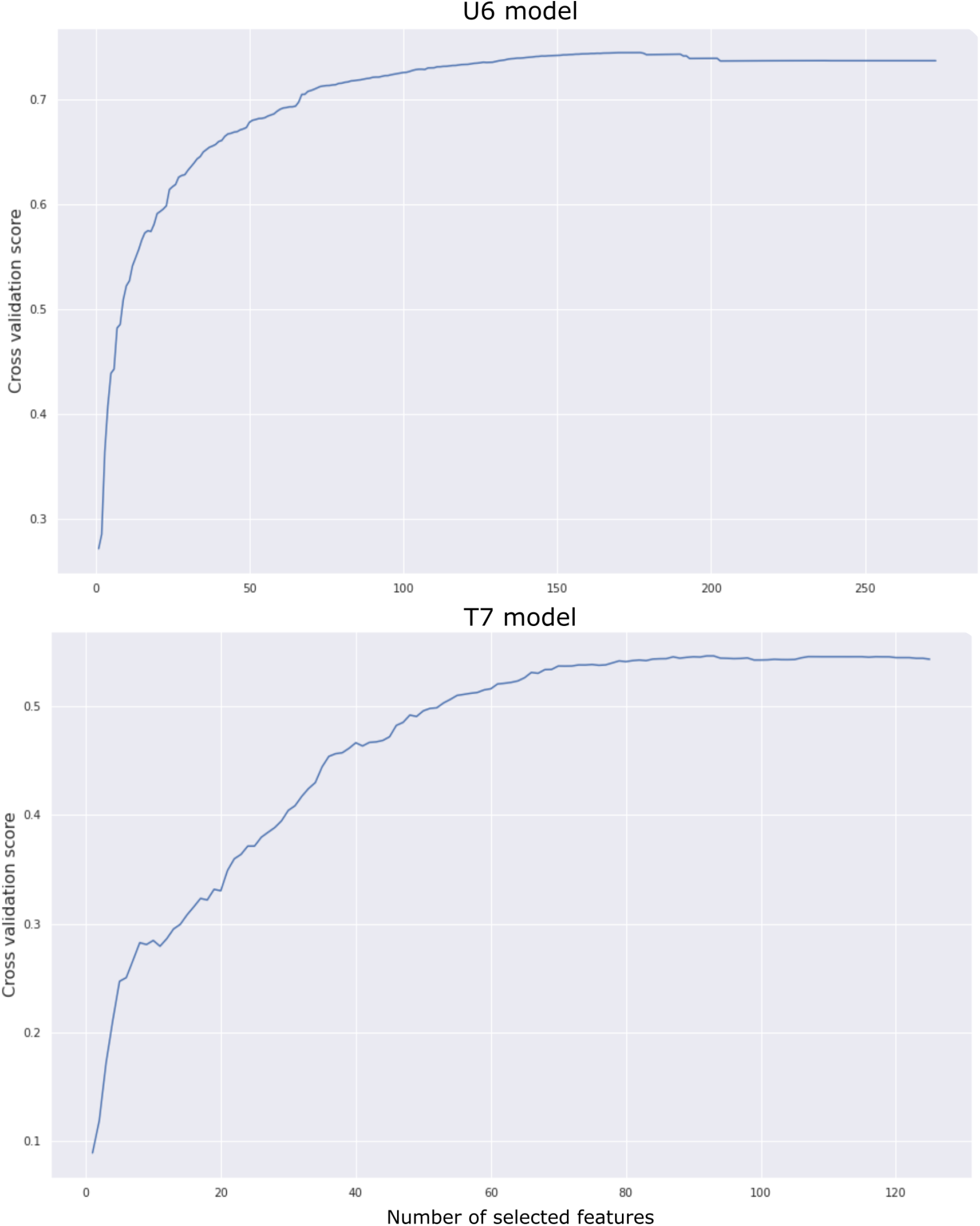
Recursive feature elimination with cross-validation for both model variants. The x-axis displays the number of selected features, while the y-axis presents the resulting performance based on Spearman correlation.

We compared five different algorithms after each step of feature selection using 10-fold cross-validation. These algorithms, described in Section 2.2.3, were evaluated in order to determine the appropriate one for our task. The results of this comparison are shown in Figure 2. Regarding the U6 dataset, we noticed that LR was inferior only to SVR for a small margin. Moreover, LR outperformed every algorithm on the T7 dataset. Based on these results and the goal of our study, we chose the model with the highest performance and lowest complexity. Therefore, Linear Regression - which was the most accurate and interpretable algorithm - was the model of our choice. Finally, we confirmed that the feature selection strategy was successful. It substantially reduced the number of features from 2466 to 28/25, while keeping an equivalent - if not better - performance.

**Figure 2:**
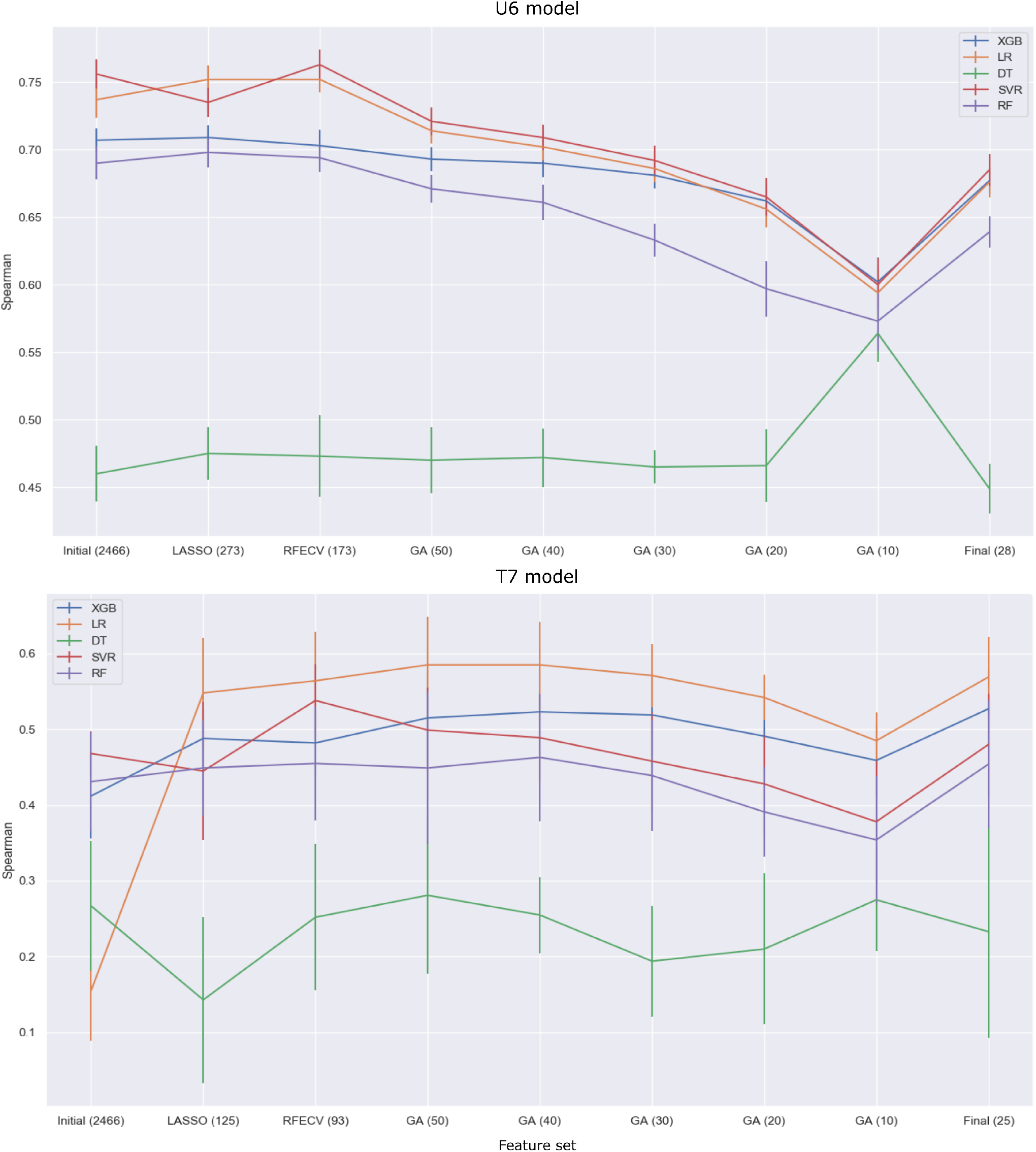
Performance comparison of feature selection strategies under 10-fold cross-validation. The number of features for each set is given in parentheses. LASSO: least absolute shrinkage and selection operator, RFECV: recursive feature elimination with cross-validation, GA: genetic algorithm.

After determining the final features and algorithms, we constructed one regression and one classification model for each dataset. In particular, we trained a binomial regression and a linear regression model to predict the percentage of efficient edits for the U6 and T7 variant. Moreover, we used the same (i.e., 28 and 25) features to train two logistic regression models by labeling the top 20% and bottom 20% gRNAs as efficient and inefficient. Therefore, we created 4 models that can be used for various tasks and under different experimental conditions. All models were implemented using the Statsmodels [43] library in Python.

### 2.3 State-of-the-art tools

We compared *CRISPRedict* to eight state-of-the-art gRNA design tools, including three machine learning (i.e., *Azimuth 2*.*0, TSAM, sgDesigner*) and five deep learning ones (i.e., *DeepCRISPR, DeepCas9, DeepSpCas9, DeepHF, CRISPRLearner*). *DeepCRISPR* [19] predictions were generated using the command-line version with sequence features only. The source R code of *DeepCas9* [20] was downloaded from https://github.com/lje00006/DeepCas9 and used with the provided weights. *DeepSpCas9* [21] and *DeepHF* [22] predictions were computed using the provided source codes available at https://github.com/MyungjaeSong/Paired-Library/tree/DeepCRISPR.info/DeepCas9 and https://github.com/izhangcd/DeepHF, respectively. Regarding *CRISPRLearner* [23], we selected the appropriate model for each dataset. We used the “Doench mEL4”, “Chari 293T”, and “Moreno-Mateos Zebrafish” models without any changes. However, due to reproducibility issues,

we retrained the “Wang-Xu HL60” and “Doench Hg19” models. Table 1 shows the models that were used and the respective testing datasets. Azimuth 2.0 [15] predictions were retrieved using the source code available at https://github.com/microsoftResearch/azimuth. We also used the python version of TSAM [27] for our evaluation. Similar to the case of CRISPRLearner, we selected the optimal model for each dataset. The “TSAM T7” model was used for datasets containing gRNAs that were expressed with the T7 promoter (i.e., Shkumatava, Gagnon, Varshney, Teboul [24]). We evaluated the remaining datasets with the “TSAM_U6” model. Finally, sgDesigner [25] predictions were generated using the command-line implementation. We processed the results into a suitable format and integrated all the predictions into one file for further analysis.

**Table 1:**
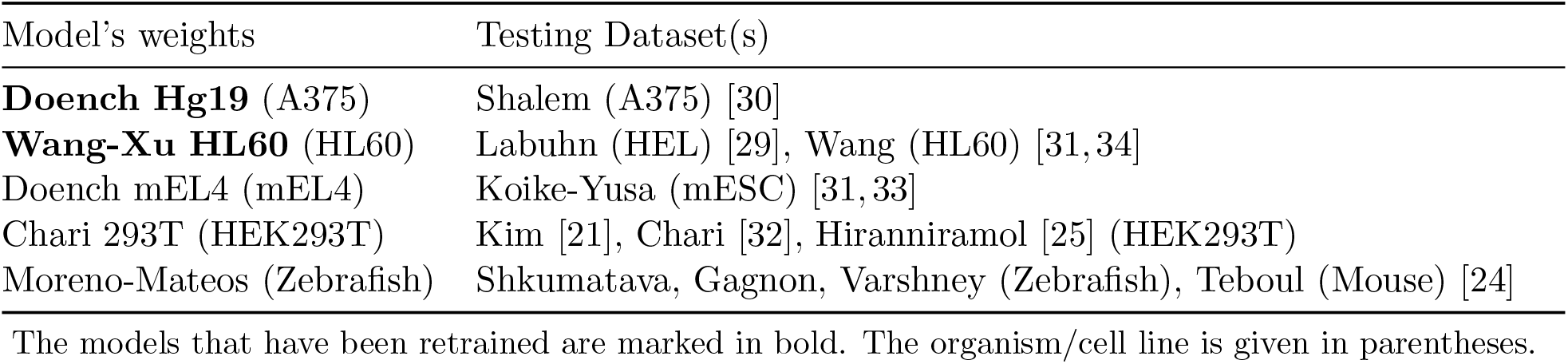
Selected *CRISPRLearner* models.

### 2.4 Evaluation metrics

Spearman correlation, normalized discounted cumulative gain (nDCG), and R-Precision were used to measure the consistency between experimentally determined gRNA efficiencies and predicted scores in the regression scenario. R-Precision and Area Under the Curve (AUC) were used for the evaluation in the classification scenario.

First, we used Spearman correlation because it does not carry any assumptions about the distribution of the data and is more robust to outliers compared with Pearson correlation [44]. It was also adopted in previous gRNA activity prediction studies [15, 18–21, 23, 24].

We applied the second metric, nDCG, to measure the ranking ability of gRNA design tools [45]. One interesting characteristic of nDCG is that the top results get more attention than the last ones through a discount function. This function can be set to zero for a specific cut-off *k*, whereby the remaining results after the *k*-th one are completely ignored [45]. This is interesting because we do not want to base our judgment on how well a tool is doing in predicting inefficiency. In our experiments, we set k to be a constant proportion of the total samples n of each dataset (i.e., *k* = *n/*5). This choice was based on the results of a previous study [45], which showed that nDCG with such cut-off converges and consistently distinguishes between various ranking functions. The nDCG value for each tool was calculated as follows: We obtained a gRNA ranking based on the predictions of the tested tool, where variable *i* represents the *i*-th gRNA in the rank list. Then, a variable *rel*_*i*_ was used to represent the relevance of the *i*-th gRNA, where *rel*_*i*_ was the real efficiency score of the corresponding gRNA on the test dataset. Finally, the performance of the tool, when considering the top *k* gRNAs, was measured by *nDCG*_*k*_ using the following formula:

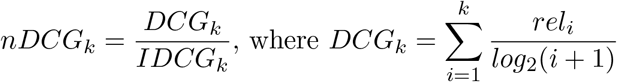

is the discounted cumulative gain (*DCG*_*k*_) for the obtained rank list and *IDCG*_*k*_ is the ideal discounted cumulative gain (i.e., a perfect ranking algorithm has *nDCG*_*k*_ = 1.0).

We also implemented the R-Precision measure [46] to evaluate the retrieval of the most efficient guide RNAs. R-precision is identical to the break-even point, where precision and recall are equal. It is also highly correlated to mean average precision [46]. Thus, it is an appropriate and informative metric for our evaluation. It is calculated as follows: Given a query (dataset) with R relevant documents (gRNAs), consider the list of documents (gRNAs) returned by a tool, and let *tot rel*(*i*) be the total number of relevant documents (gRNAs) retrieved up to and including rank *i*. The R-precision of this list is the precision at rank R, i.e.,

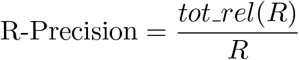

R-Precision was used for both evaluation scenarios. The number of relevant (i.e., efficient) gRNAs was determined differently in each scenario. In the regression scheme, the 80-th percentile of guide efficiencies per dataset was used as a threshold. Every gRNA with efficiency greater than or equal to this threshold was considered relevant. In the case of classification, all gRNAs that were labeled as efficient were considered relevant.

Finally, AUC of the Receiver Operating Characteristic (ROC) was used to measure the classification ability of each model [47]. The ROC curve is a plot of the True Positive Rate (TPR) versus the False Positive Rate (FPR) across different thresholds. Thus, it offers a threshold-independent way of evaluating information retrieval performance. It also introduces the AUC measure, which is a simple scalar metric that defines how an algorithm performs over the whole space.

Spearman correlation was calculated using the SciPy [48] library while nDCG and R-Precision were calculated with our custom Python scripts using the aforementioned equations. ROC analysis and AUC calculation were performed using the Scikit-learn package [40].

### 2.5 Statistical significance

To confirm that *CRISPRedict* has comparable performance to the most accurate tools, we used the two one-sided test (TOST) procedure [49]. The goal of this procedure is to determine whether two normally distributed samples come from populations that are sufficiently similar in means. As a test for equivalence, it essentially reverses the roles of the null and alternative hypothesis. Specifically, TOST assumes a null hypothesis in which the means of two distributions are not equivalent. The alternative hypothesis is that the two means are equivalent within an acceptable margin delta (*δ*). Formally, if Δ = *µ*_1_ *− µ*_2_ is the difference between the means of the two variables and *δ* is the equivalence margin, the null and alternative hypotheses are:

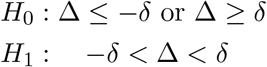

The null hypothesis is made up of two simple one-sided hypotheses:

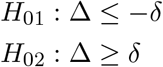

If both of these one-sided tests are rejected, we conclude that the paired variables are equivalent (i.e., their average difference is confined within a small predetermined margin). The *δ* is set by the researcher and is defined as the highest tolerated effect size difference. It allows researchers to look at equivalence between means within a small range as opposed to looking for equality, which implies two distributions that are identical in every way. We set *δ* to be equal to 0.075 for a total margin of 0.15. The level of statistical significance was set to 0.05. The normality assumption of the TOST procedure was tested using the Shapiro-Wilk test [50]. Both procedures were performed using the Pingouin package in Python [51].

### 2.6 Code and data availability

The source code of *CRISPRedict*, along with data and Python scripts to reproduce our results, is available at https://github.com/VKonstantakos/CRISPRedict. The results are also available in the Supplementary material (Table S1).

## 3 Results

To assess its general performance, we evaluated *CRISPRedict* using published datasets, derived from studies from independent laboratories. These datasets, described in Section 2.1.2, include gRNAs expressed with the U6 or the T7 promoter and represent the guide efficiency using a continuous or a discrete variable, corresponding to the regression and classification tasks supported by *CRISPRe-dict*. Therefore, they allow to evaluate our tool under different experimental conditions and tasks which are grouped into two testing scenarios, one for regression and one for classification. For a fair comparison, we curated data which were not used for training and excluded the results of models tested against their own training datasets. With these datasets, we compared *CRISPRedict* with the current state-of-the-art gRNA design tools, including *DeepCRISPR* [19], *DeepCas9* [20], *DeepSpCas9* [21], *DeepHF* [22], *CRISPRLearner* [23], *Azimuth 2*.*0* [15], *TSAM* [27], *and sgDesigner* [25]. We also applied equivalence testing to confirm that *CRISPRedict* has comparable performance to the most accurate tools, as shown by our evaluation. Finally, we visualized *CRISPRedict* ‘s predictions in order to demonstrate the additional interpretability of our implementation.

### 3.1 Comparison of CRISPRedict with state-of-the-art gRNA design tools

#### 3.1.1 Testing scenario 1 - Regression scheme

In this section, we present the performance of *CRISPRedict* under a regression scheme. Specifically, we measure the consistency between experimentally validated gRNA efficiencies and predicted scores on eight independent datasets, using Spearman correlation, normalized discounted cumulative gain (nDCG), and R-Precision. The results of this evaluation can be seen in Figure 3 and 4. We observe that *CRISPRedict* performs well, among all datasets and metrics. In particular, it is consistently among the three most accurate models, as shown in Figure 4. Moreover, it has the most stable performance for both U6 and T7 datasets. Similarly, good and stable performance is evident in the case of *DeepHF* [22] *and TSAM* [27], which also used distinct model variants for the U6 and T7 datasets. On the other hand, DeepSpCas9 [21], a tool with high generalization performance [18], is the most accurate on the U6 datasets but demonstrates a decreased and variable performance on the T7 datasets. Finally, sgDesigner [25], a stacking framework trained on a plasmid library under a U6 promoter, has a stable, albeit lower, accuracy for both U6 and T7 data.

**Figure 3:**
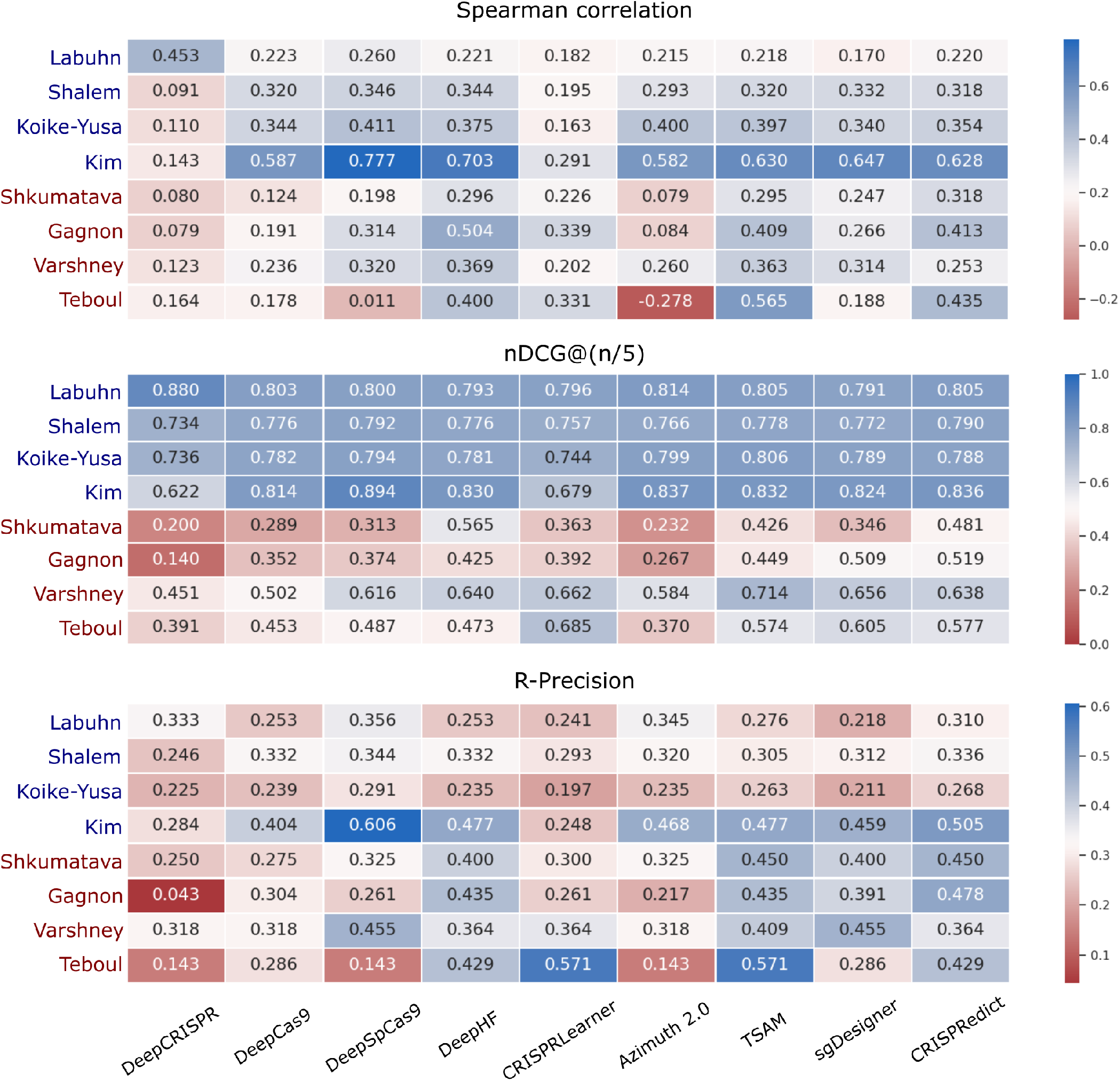
Evaluation of the *CRISPRedict* regression model. The heat maps show the performance of *CRISPRedict* and previously reported models, when assessed with Spearman correlation, nDCG, and R-Precision. Each row represents a testing dataset, while each column represents an evaluated model. Testing datasets that include gRNAs expressed from a U6 promoter or a T7 promoter are colored blue and red, respectively. nDCG: normalized discounted cumulative gain.

**Figure 4:**
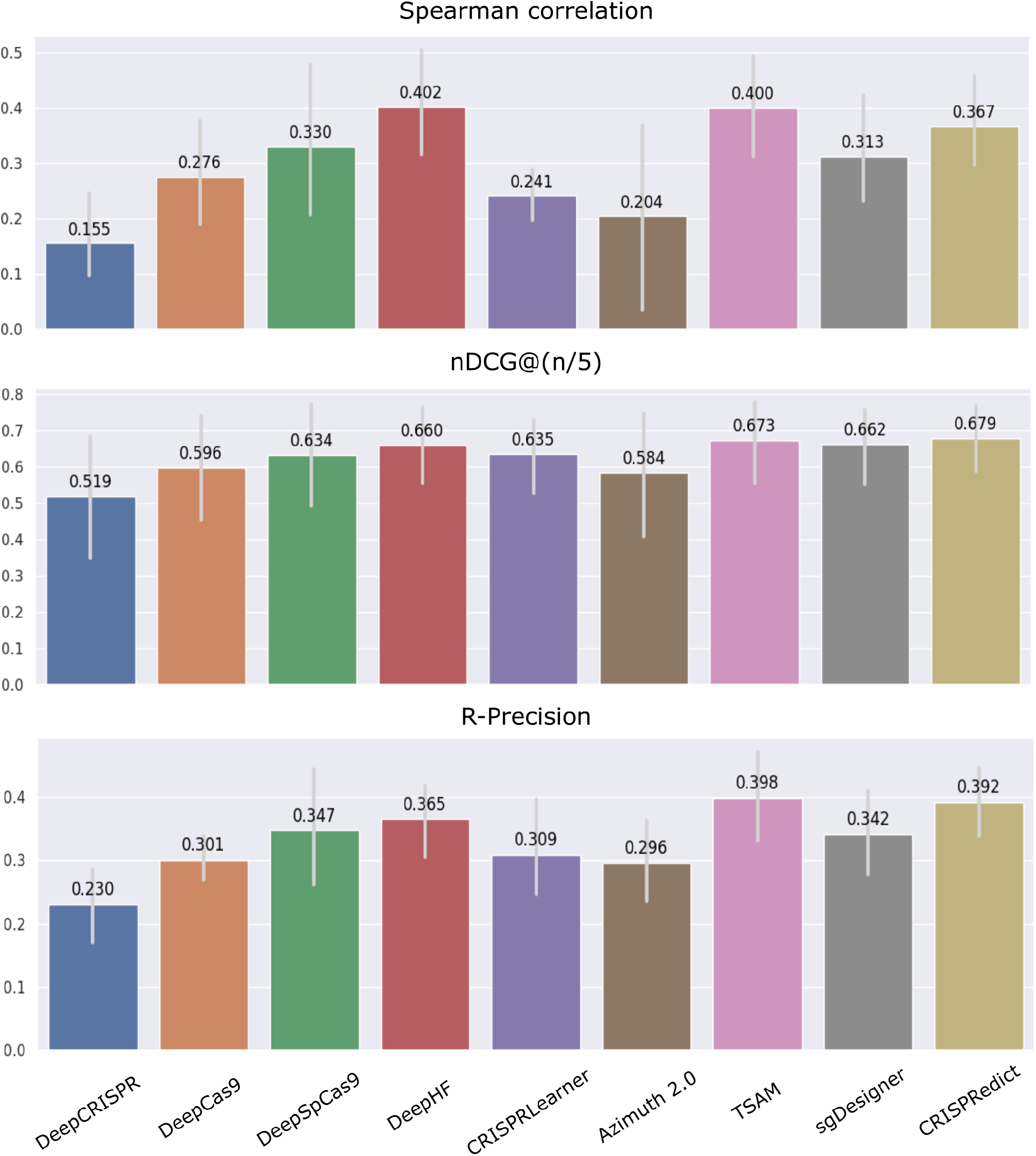
Comparison of the general performance of the *CRISPRedict* regression model. Each bar demonstrates the mean performance of a model across all eight datasets, using a single evaluation metric. nDCG: normalized discounted cumulative gain.

The remaining tools perform worse than the aforementioned ones, especially DeepCRISPR [19], which has the best performance in the HEL cell line but performs poorly in the other datasets. Thus, we conclude that CRISPRedict demonstrates a good and robust performance, despite being a simple linear model.

#### 3.1.2 Testing scenario 2 - Classification scheme

Although our main goal was to provide a model that can accurately rank potential target sites, we also created a classifier to label the candidate gRNAs as efficient or inefficient. Since most of the existing tools only provide a regression model, *CRISPRedict* can fill this gap and help researchers differentiate between highly active and less active gRNAs without requesting an accurate prediction of the efficiency value itself.

In this section, we assess *CRISPRedict* ‘s classification ability on four independent datasets. These datasets include labeled gRNAs expressed under a U6 promoter. To compare all the current state-of-the-art tools, including regression and classification ones, we used R-Precision and Area Under the Curve (AUC). The results of this comparison are shown in Figure 5.

**Figure 5:**
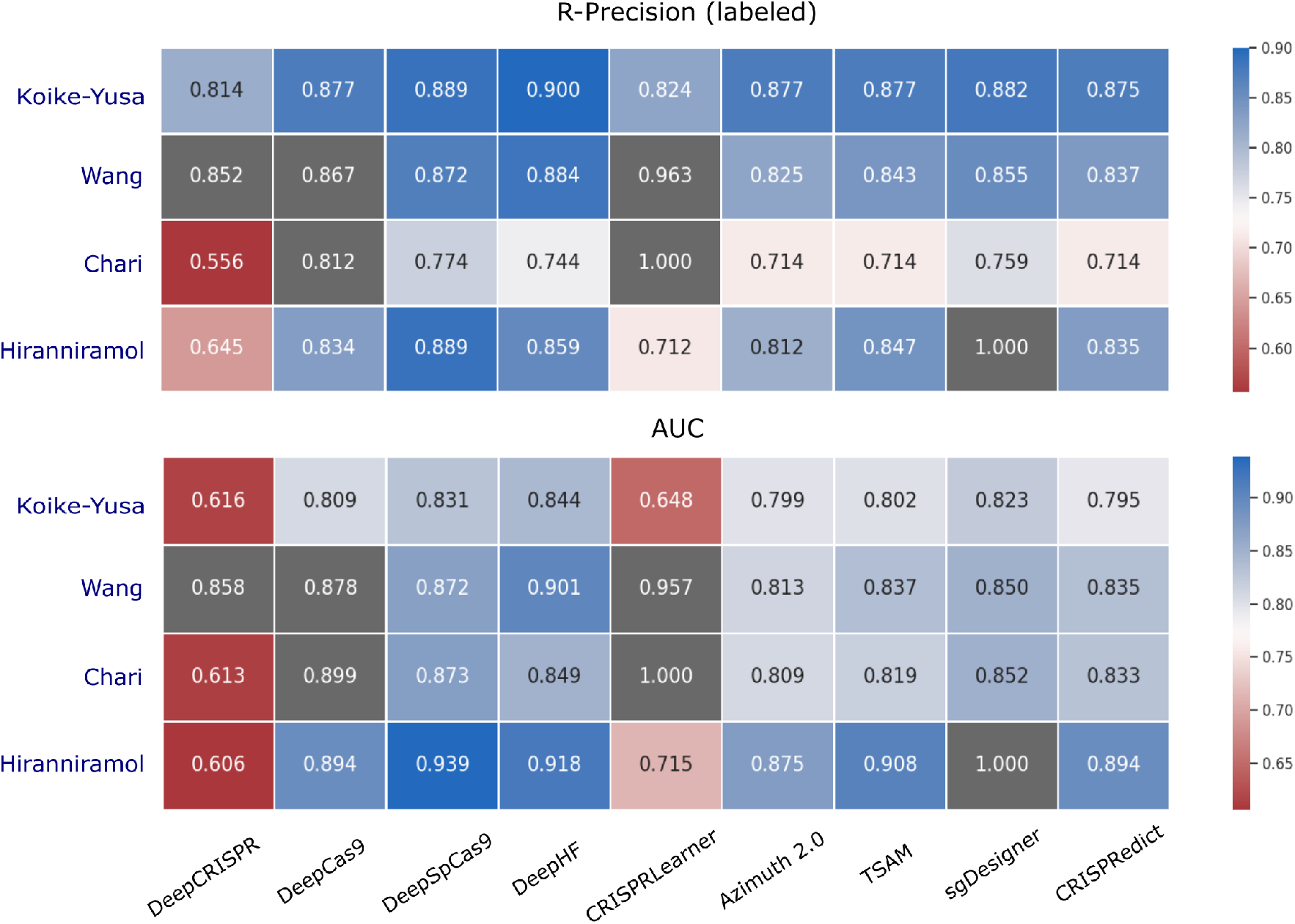
Evaluation of the *CRISPRedict* classification model. The heat maps show the performance of *CRISPRedict* and previously reported models, when assessed with R-Precision and AUC. Each row represents a testing dataset, while each column represents an evaluated model. Testing datasets that include gRNAs expressed from a U6 promoter are colored blue. Evaluations of an algorithm against its own training data set are shown in gray as they are likely to be overestimated due to over-fitting. AUC: area under the curve.

First, we observe that the performances of all models are generally stable since the testing datasets include gRNAs expressed only from a U6 promoter. In addition, all the models demonstrate comparable performances, possibly due to the nature of the problem. In particular, classification, being less informative, is an easier task than predicting the exact efficiency of a gRNA (regression). Therefore, the differences between the models become less evident and the comparison is more difficult. For this reason, the bar charts and the Receiver Operating Characteristic (ROC) curves are not helpful in this case. Having said that, the models with the highest classification performance are *DeepSpCas9* and *DeepHF*, while *CRISPRedict* ‘s accuracy is on a similar but slightly lower level. However, we argue that these differences are not statistically significant and *CRISPRedict* provides the additional benefit of a clear threshold for labeling gRNAs. We study this hypothesis in the following section.

#### 3.1.3 Statistical testing

To test whether *CRISPRedict* has comparable performance to the most accurate tools, we implemented an equivalence testing approach. In particular, we used the two one-sided test (TOST) procedure [49] to compare *CRISPRedict* with the top-performing machine learning and deep learning tool in our evaluation, namely *TSAM* and *DeepHF*. This way, we can confirm that our tool is at least as accurate as the current state-of-the-art tools with the additional benefit of being interpretable.

The TOST procedure assumes a null hypothesis in which the performances of the two compared models are not equivalent (Section 2.5). The first model is either better or worse than the second by a margin of delta (*δ*). If both of these hypotheses are rejected, the alternative hypothesis holds. Thus, the models’ performances are equivalent within an acceptable difference *δ*. We used this test to compare *CRISPRedict* with *TSAM* and *DeepHF* for all the described metrics. The results of the statistical procedures are shown in Table 2.

**Table 2:**
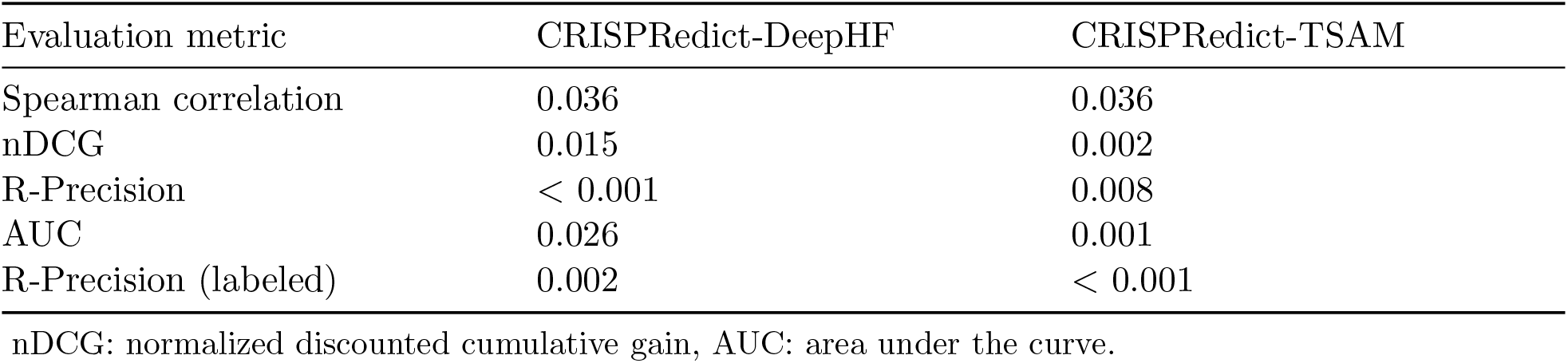
P-values of the TOST procedure between CRISPRedict and DeepHF/TSAM.

We observe that all the resulting p-values are less than 0.05, obtaining statistically significant results for both model comparisons. This illustrates that *CRISPRedict* has equivalent performance to *TSAM* and *DeepHF* when evaluated with correlation, ranking, and classification metrics. Despite its simplicity, *CRISPRedict* can compete with the most accurate tools and even outperform some of them.

### 3.2 Visualizing and interpreting CRISPRedict’s prediction

Previous studies have addressed the problem of feature identification and visualization using conventional machine learning and deep learning methods [15, 19, 20, 22, 30, 31, 52, 53]. In the first case, feature importance weights and statistical tests have been used to identify features that significantly affect the cleavage efficiency. In the case of deep learning tools, various approaches such as feature salience maps [19], SHAP values [22], and permutation methods [20] have been implemented in order to interpret their predictions.

However, both approaches have limitations. First, the implemented tools are still black-box models that do not provide intrinsically explainable predictions. Moreover, the described methods for interpretability give mostly global information about the model and not about individual predictions. In the case of machine learning models, the feature engineering process may lead to many redundant features that further hinder their explainability. For example, *Azimuth 2*.*0* [15] and sgDesigner [25], include 582 and 302 features, respectively, making their predictions hard to explain. Similarly, the deep learning tools are also difficult to interpret and sometimes lead to conflicting results about the important gRNA features.

In this section, we highlight the ability of *CRISPRedict* to directly provide interpretable and explainable predictions for the CRISPR-Cas9 system. To accomplish that, we use two types of visualizations: (i) color-based nomograms [54] and (ii) effect plots [55]. These visualizations can provide both general information about the model and specific explanations about an instance. An example of each plot is shown in Figure 6 and 7, respectively.

**Figure 6:**
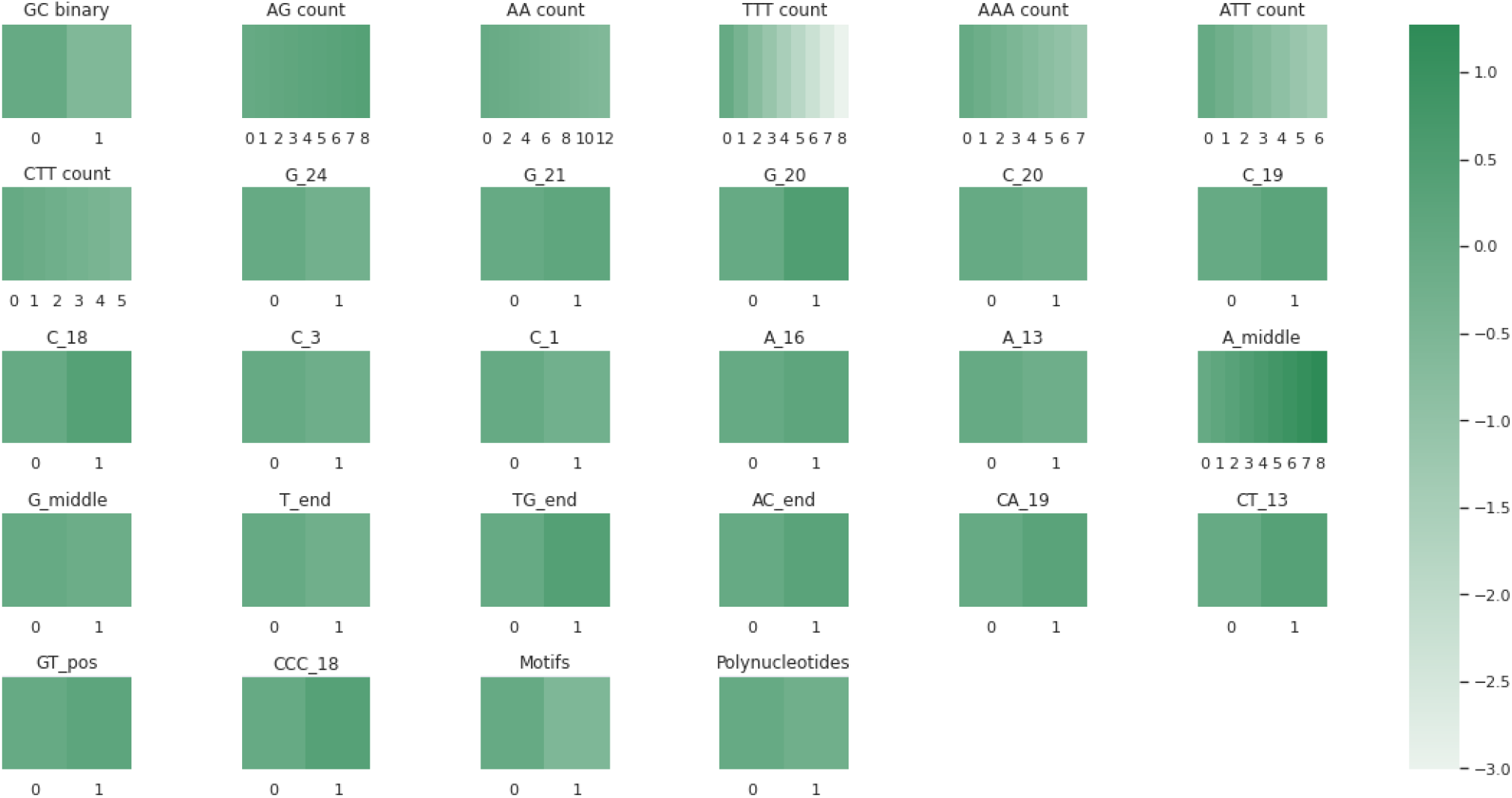
Visualizing the contribution of each feature with a color-based nomogram.

**Figure 7:**
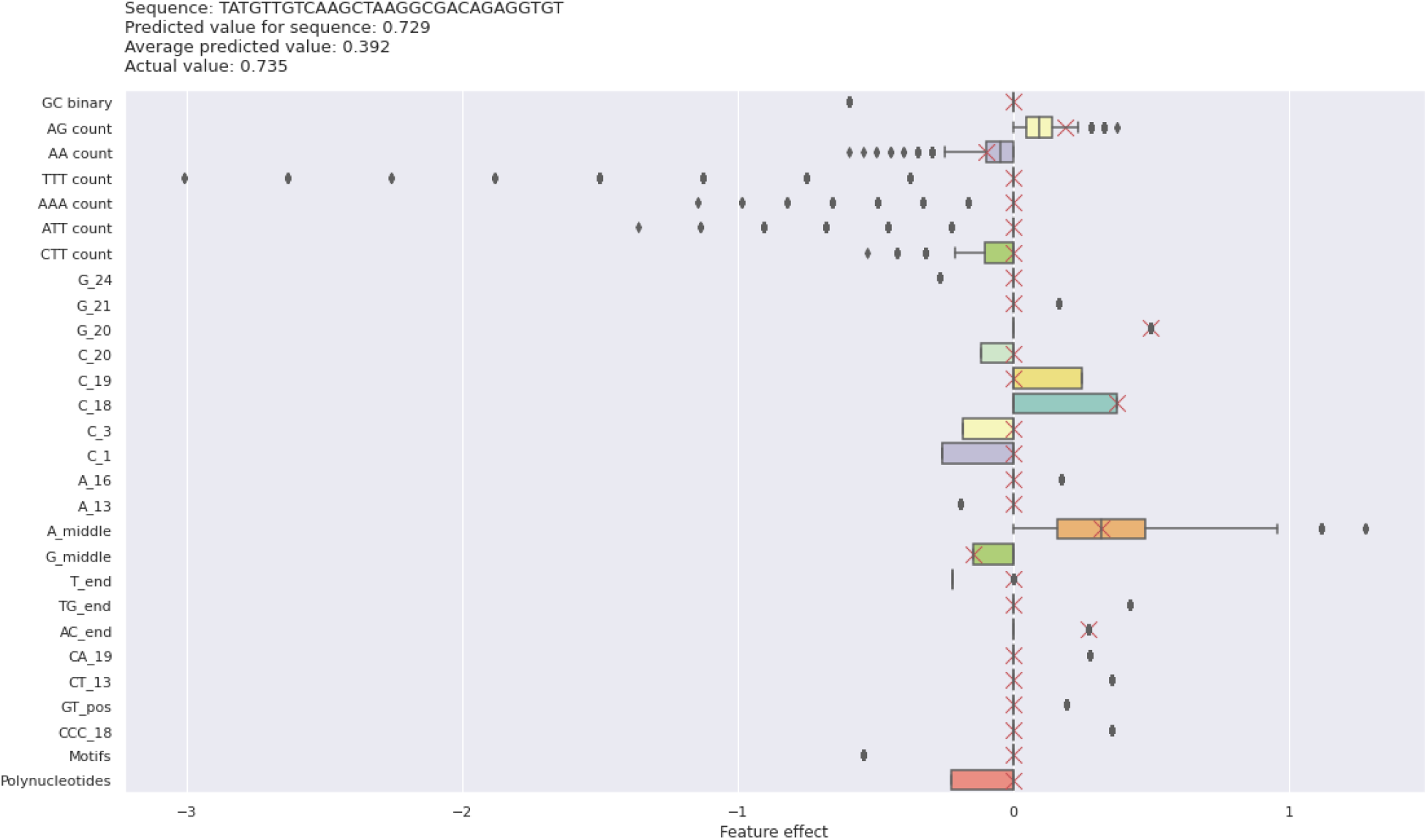
Explaining model’s predictions with an effect plot.

Borrowing from risk prediction models [54, 56], the color-based nomogram illustrates the contribution of each feature to the final prediction. For instance, we observe that the presence of G in position 20 and the TTT count are significant predicting variables in the provided example (Fig. 6). In addition, we can directly obtain instance-based explanations by marking the feature values of a sequence and calculating the sum of all their contributions (Supplementary Fig. S1).

Similarly, the weights of a linear model can be more meaningfully analyzed when they are multiplied by the actual feature values. Moreover, it is important to know the distribution of the features in the data, because if there is very low variance, almost all instances will have similar contribution from this feature. The effect plot [55] can help understand how much the combination of weight and feature contributes to the final predictions. It can also be used to explain individual predictions by marking the actual feature value of an instance and comparing it with the general feature distribution (Fig. 7).

## 4 Conclusion

The CRISPR-Cas9 system has rapidly emerged as a state-of-the-art technology for genome editing applications. Because of its simplicity, efficacy, specificity, and versatility, this technology has tremendous advantages over other gene-editing technologies. Current gRNA design tools serve as an important platform for the efficient application and development of the CRISPR system. However, the existing models still have some limitations, such as their variable accuracy and unclear mechanism of decision making.

In this study, we address these issues by introducing a simple and interpretable linear model, named *CRISPRedict*, for gRNA efficiency prediction. After a comprehensive evaluation, we demonstrate that *CRISPRedict* has an equivalent performance with the currently most accurate tools and outperforms the remaining ones. Moreover, it has the most robust performance for both U6 and T7 datasets, illustrating the general applicability of our model to tasks under different experimental conditions. Finally, since most of the existing tools provide only regression models, we also create a classification variant of *CRISPRedict* that provides the additional benefit of a clear threshold for labeling gRNAs. Given the performance, interpretability, and versatility of *CRISPRedict*, we expect that it will greatly facilitate genome editing using CRISPR-Cas9.

