## Supplementary Figure 1 for "CRISPRedict: The case for simple and interpretable efficiency prediction for CRISPR-Cas9 gene editing"

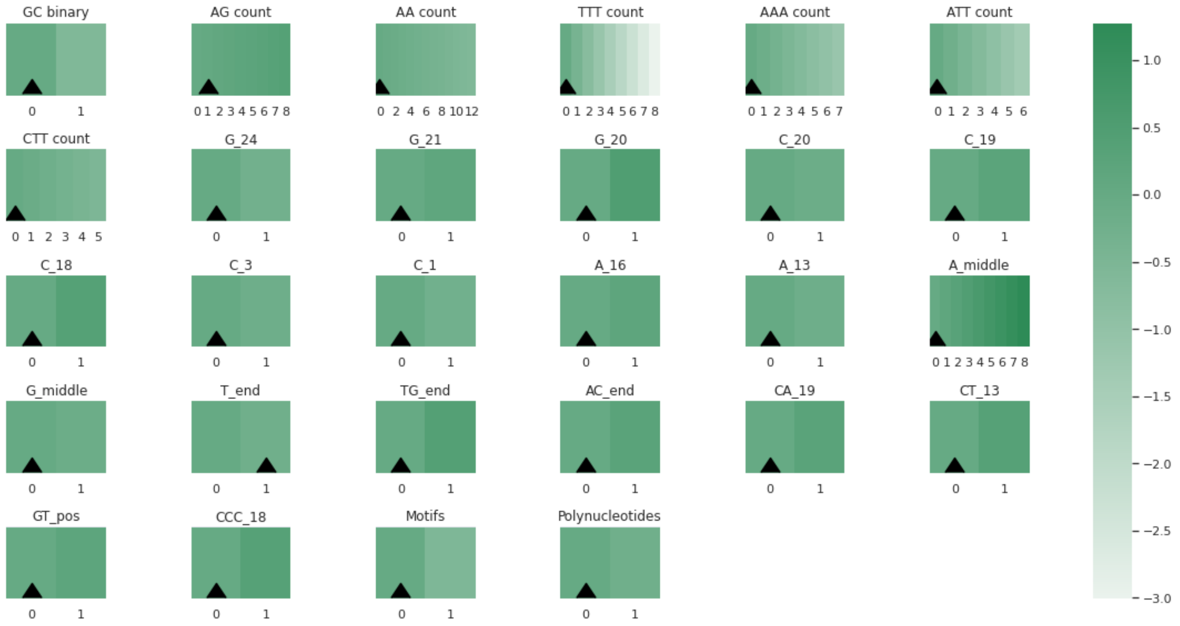


Supplementary Figure 1: Visualizing the contribution of each feature with a color-based nomogram. The feature values of the provided sequence have been marked to obtain an instance-based explanation.
